## Supplemental Information for "Interplay of structured and random interactions in complex ecosystems dynamics"

### Supporting Information for "Interplay of structured and random interactions in complex ecosystems dynamics"

#### A Derivation of the dimensionless equations

Let us consider a community of $S$ interacting species labeled $i = 1, \ldots, S$, of respective abundances $N_i(t)$. The Lotka-Volterra equations are the simplest non-linear model capturing the effect of ecosystem composition on individual demographic rates [1]. They reproduce qualitative features of population dynamics, such as exponential growth or decay, stable survival and extinction. Within this framework, each species is endowed with an intrinsic net growth rate $r_i$. Species with larger $r_i$ have faster dynamics, and therefore react faster to changes in community composition. The species' carrying capacity $K_i$ is the maximum abundance it can attain in isolation as a result of competition between individuals of the same species.

Interactions with other species are characterized by a matrix of interaction coefficients $a_{ij}$ that quantify the impact of species $j$ on the instantaneous growth rate of species $i$. In our convention, a negative $a_{ij}$ implies a deleterious effect on growth and thus competition; on the contrary, a positive coefficient indicates a facilitative interaction. The abundance of every species then changes in time according to

$$\frac{\mathrm{d}N_i}{\mathrm{d}t} = r_i N_i \left[1 - \frac{N_i - \sum_j a_{ij} N_j}{K_i}\right] + m_i, \tag{S1}$$

Migration into the community has also been added through the *immigration rates* $m_i$ [2, 3]. In numerical simulations, $m_i \ll K_i$. Some species that persist at the immigration level can therefore invade as soon as the community context becomes favorable. At any given time, we count as extinct those species whose abundance is smaller than a multiple of $m_i$.

In order to focus on the effect of variability in the interaction parameters, we make the simplifying assumption that some parameters are uniform across species and rescale eq. (S1) so that variables become independent of the carrying capacity. The analytical tools however generalize to cases when this transformation is not realized. Assuming that $r_i$ is constant across species, it can be set to one, without loss of generality, by rescaling the time coordinate. We also set $m_i = m$ for all species. Finally, by factoring out $K_i$, one can rewrite the Generalized Lotka-Volterra Equations (GLVs) in terms of the normalized abundances $x_i = N_i/K_i$ and the rescaled interaction coefficients $A_{ij} = a_{ij}K_j/K_i$ as

$$\frac{\mathrm{d}x_i}{\mathrm{d}t} = x_i \left[1 - x_i + \sum_j A_{ij} x_j\right] + m. \tag{S2}$$

### B Properties of the interaction matrix

#### B.1 Scaling

We aim to work in a regime where ecological processes stem from aggregating the contribution from many different species. Given the dynamical equation

$$\frac{\mathrm{d}x_i}{\mathrm{d}t} = x_i \left(1 - x_i + \sum_j A_{ij} x_j\right), \tag{S3}$$

this means that the term $A\vec{x}$ needs to be $\mathcal{O}(1)$ even in the presence of an extensive number of species that all have abundances $\mathcal{O}(1)$. Given that

$$\mathbb{E}\left[\sum_j A_{ij}x_j\right] \sim \sum_j \mathbb{E}\left[A_{ij}\right] \sim S\mathbb{E}\left[A_{ij}\right] \tag{S4}$$

$$\mathbb{V}\text{ar}\left[\sum_j A_{ij}x_j\right] \sim \sum_j \mathbb{V}\text{ar}\left[A_{ij}\right] \sim S\mathbb{V}\text{ar}\left[A_{ij}\right] \tag{S5}$$

we need both the mean and the variance of the interaction coefficients to go as $1/S$. This leads to the decomposition of the main text,

$$A_{ij} = \mu_{ij} + \frac{\sigma}{\sqrt{S}} z_{ij}, \tag{S6}$$

where $\mu_{ij} \sim 1/S$.

#### B.2 Rank and eigenvectors of the structure matrix

For the reasons stated in the main text, the structure matrix is assumed to be low-rank. Although our main justification is based on the notion of macroscopic structure, low-rank is also an automatic consequence of the scaling chosen in eq. (S6). Indeed, we have

$$\text{Tr}\left[\mu^T \mu\right] = \sum_{ij} \mu_{ij}^2 \sim \mathcal{O}(1). \tag{S7}$$

On the other hand this trace is also the sum of the singular values of $\mu$, therefore only a finite number of them can be of order 1, i.e. $\mu$ is low-rank. Now we turn to the eigenvectors of $\mu$. First, let $\vec{x}$ be a localized vector. Then

$$\left[\mu\vec{x}\right]_i = \sum_{j:x_j\neq 0} \mu_{ij}x_j \ll 1. \tag{S8}$$

Therefore the structure part doesn't act on localized directions. Similarly, assume that the vector is sampled from a uniform distribution on the unit sphere

$$\|\mu\vec{x}\|^2 = \sum_{ij} x_i\mu_{ki}\mu_{kj}x_j = \mathcal{O}(S^{-2}). \tag{S9}$$

Therefore neither does it act on 'typically random' directions. In particular, eigenvectors of $\mu$ with non-zero eigenvalue are collective and need to have non-random features in their entries (for instance, having more positive than negative entries, or be structured in groups).

Conversely, assume that the matrix is low-rank and write it using its Singular Value Decomposition

$$\mu_{ij} = S^{-1} \sum_{\lambda} \mathcal{S}_i^{(\lambda)} \mathcal{I}_j^{(\lambda)}, \tag{S10}$$

where the $\lambda$'s are the (positive) singular values, which are $\mathcal{O}(1)$ and the $\mathcal{S}^{(\lambda)}$ and $\mathcal{I}^{(\lambda)}$ are non-normalized. Then given that $\mathcal{I}_j^{(\lambda)}, \mathcal{S}_i^{(\lambda)} \sim \mathcal{O}(1)$, the scaling of the interaction matrix is $\mathcal{O}(S^{-1})$. Furthermore, if the eigenvectors were randomly distributed then the matrix would only act on the very low-dimensional space spanned by those vectors. In particular, the projection of any fixed vector with positive entries on this low-dimensional is almost surely zero. Given that the interaction matrix is applied to abundance vectors, which have positive entries, such matrices would effectively imply vanishing interactions in the Lotka-Volterra model. Hence eigenvectors of low-rank matrices need to be non-random for them to be meaningful.

Therefore we see that there is a quantitative equivalence between the $1/S$ scaling for the structural part and the fact of having a low rank and non-random eigenvectors. In other words, collective structure and this scaling go hand-in-hand, therefore providing a mathematical justification for our choice. This is the complete opposite of what happens in the random part of the interactions, where the scaling can be upgraded to $1/\sqrt{S}$ precisely because the eigenvectors are random and the rank is full.

### C Numerical simulations and solutions

#### C.1 Numerical simulations

All numerical calcualtions where performed in Python. Numerical simulations where done using Scipy's `solve_ivp` function with the `RK45` method. Interactions for simulations shown in the main text were drawn from Gaussian distributions although other distributions were also tested. Initial conditions were sampled uniformly between zero and 1. The solver was stopped after a fixed amount of time, generally $T = 1000$. This is more than enough for convergence in the UFP phase and also enough to observe a large-enough amount of cycles in the cyclic phase. The immigration threshold was set to $\varepsilon = 10^{-8}$.

##### C.2 Numerical solutions

Numerical solutions for the fixed point equations where obtained by using an iterative scheme. The equations where rewritten in vector form as $\vec{X} = f(\vec{X})$ and the iteration ran as $\vec{X}_{n+1} = (1-r)\vec{X}_n + rf(\vec{X}_n)$ for a fixed number of iterations. We used $r = 0.25$. The initial condition was taken to be $\vec{X}_0 = \vec{1}$. Although we have no mathematical guarantee of convergence for this solver beyond the unstructured case, we find that it to converge in practice as long as a solution exists. Performance-wise, the solver runs extremely fast due to the low-dimensionality of the space on which the equations are rewritten. Except very close to the divergence transition, we find that 200 iterations were more than enough to converge to the solution, although the examples of the main text were solved with 20000 iterations.

For the examples of the article, this simple scheme was enough. For more complicated cases, several solutions to the fixed point equations can exist, corresponding to different structural regimes. In that case, the initial condition plays a role in determining towards which solution the solver converges. For those cases, we used a combination of multiple initial conditions and intuitive guessing to find the solutions, but a more automatic solver should be implemented for future work.

#### D Obtaining the effective process with Dynamical Mean Field Theory

We now describe how to go from the deterministic Lotka-Volterra equations with random prescription for the interaction matrix to the set of stochastic single-species processes with dynamical noise. Recalling that the trajectories of single species are random due to the randomness in the interactions the idea is to express their probability distribution and simplify it to a point where it can be recognized as the probability density associated to a stochastic process. Start by writing the probability of a trajectory $\vec{x}(t)$ as

$$\mathbb{P}\left[\vec{x}(t)\right] = \int \prod_{ij} \mathrm{d}z_{ij} \mathbb{P}\left[z_{ij}\right] \prod_i \delta_f \left( \frac{\dot{x}_i}{x_i} - 1 + x_i - \sum_j \left( \mu_{ij} + \frac{\sigma}{\sqrt{S}} z_{ij} \right) x_j \right), \tag{S11}$$

where $\delta_f$ stands for a functional Dirac delta function, enforcing the dynamics at each time step. Indeed, the equation inside the Delta is equivalent to the gLV equation for the $i^{\text{th}}$ species, provided that $x_i \neq 0$. Note in particular that as long as $x_i(0) \neq 0$ or there is migration, the fact that the gLV equation is a first-order differential equation ensures that $x_i(t) \neq 0$ at any finite time. Species for which $x_i(0) = 0$ can, in the absence of migration, be removed from the community without changing the dynamics. Hence the division by $x_i$ in eq. (S11) is safe. Formally, this is just a change of variables. Seeing the functional delta as a product of deltas for each time step, we can use the Fourier representation

$$\delta\left(x\right)=\int\frac{\mathrm{d}\hat{x}}{\sqrt{2\pi}}e^{i\hat{x}x},\tag{S12}$$

so that the probability of a trajectory becomes

$$\mathbb{P}\left[\vec{x}(t)\right]=\int\mathcal{D}\hat{x}(t)\int\prod_{ij}\mathrm{d}z_{ij}\mathbb{P}\left[z_{ij}\right]\exp\left(i\sum_i\int\mathrm{d}t\hat{x}_i(t)\left(\frac{\dot{x}_i(t)}{x_i(t)}-1+x_i(t)-\sum_j\left(\mu_{ij}+\frac{\sigma}{\sqrt{S}}z_{ij}\right)x_j(t)\right)\right),\tag{S13}$$

where $\mathcal{D}\hat{x}(t)$ formally stands for a product of pointwise measures $\mathcal{D}\hat{x}(t)=\prod_t\mathrm{d}\hat{x}(t)$. In writing this and in the following, we ignore the prefactors as we will not be interested in eventual normalizing constants. With this representation we can now do the integral over the random matrix entries. Indeed,

$$\int\prod_{ij}\mathrm{d}z_{ij}\mathbb{P}\left[z_{ij}\right]\exp\left(i\sum_{ij}\int\frac{\sigma z_{ij}}{\sqrt{S}}\mathrm{d}t\hat{x}_i(t)x_j(t)\right)=\exp\left[-\frac{S\sigma^2}{2}\int\mathrm{d}t\mathrm{d}t'\left(\frac{1}{S}\sum_i\hat{x}_i(t)\hat{x}_i(t')\right)\left(\frac{1}{S}\sum_i x_i(t)x_i(t')\right)\right].\tag{S14}$$

Define the quantities

$$\mathcal{A}(t,t')=S^{-1}\sum_i x_i(t)x_i(t')\tag{S15}$$

$$\mathcal{B}(t,t')=S^{-1}\sum_i \hat{x}_i(t)\hat{x}_i(t')\tag{S16}$$

which we will later introduce in eq. (S13) using again a Fourier representation for functional deltas,

$$\int\mathcal{D}\hat{\mathcal{A}}(t,t')\mathcal{D}\mathcal{A}(t,t')\mathcal{D}\hat{\mathcal{B}}(t,t')\mathcal{D}\mathcal{B}(t,t')\exp\left(iS\int\mathrm{d}t\mathrm{d}t'\hat{\mathcal{A}}(t,t')\left(\mathcal{A}(t,t')-\frac{1}{S}\sum_i x_i(t)x_i(t')\right)\right)\tag{S17}$$

$$\times\exp\left(iS\int\mathrm{d}t\mathrm{d}t'\hat{\mathcal{B}}(t,t')\left(\mathcal{B}(t,t')-\frac{1}{S}\sum_i \hat{x}_i(t)\hat{x}_i(t')\right)\right).\tag{S18}$$

By assumption, the structural matrix $\mu_{ij}$ has low rank. We can therefore write it as

$$\mu_{ij}=S^{-1}\sum_{\lambda=1}^{\mathrm{rk}(\mu)}\mathcal{S}_i^{(\lambda)}\mathcal{I}_j^{(\lambda)}.\tag{S19}$$

This decomposition is not unique. It can stem from knowledge of the different functions whose superposition gives the matrix $\mu_{ij}$, or by a singular value decomposition. In any case, we define the following quantities

$$f_\lambda(t) = S^{-1} \sum_i \mathcal{I}_i^{(\lambda)} x_i(t) \tag{S20}$$

$$n_\lambda(t) = S^{-1} \sum_i \mathcal{S}_i^{(\lambda)} \hat{x}_i(t) \tag{S21}$$

which we introduce similarly in eq. (S13) using a Fourier representation,

$$\prod_\lambda \int \mathcal{D}\hat{f}_\lambda(t)\mathcal{D}f_\lambda(t)\mathcal{D}\hat{n}_\lambda(t)\mathcal{D}n_\lambda(t) \exp\left(iS\int \mathrm{d}t \hat{f}_\lambda(t)\left(f_\lambda(t) - \frac{1}{S}\sum_i \mathcal{I}_i^{(\lambda)} x_i(t)\right)\right) \tag{S22}$$

$$\times \exp\left(iS\int \mathrm{d}t \hat{n}_\lambda(t)\left(n_\lambda(t) - \frac{1}{S}\sum_i \mathcal{S}_i^{(\lambda)} \hat{x}_i(t)\right)\right). \tag{S23}$$

All in all the probability of a trajectory becomes

$$\mathbb{P}\left[\vec{x}(t)\right] \propto \int \mathcal{D}\mathcal{A}\mathcal{D}\hat{\mathcal{A}}\mathcal{D}\mathcal{B}\mathcal{D}\hat{\mathcal{B}} \prod_\lambda \mathcal{D}f_\lambda \mathcal{D}\hat{f}_\lambda \mathcal{D}n_\lambda \mathcal{D}\hat{n}_\lambda \tag{S24}$$

$$\times \exp\left(iS\int \mathrm{d}t\mathrm{d}t'\left(\hat{\mathcal{A}}\mathcal{A} + \hat{\mathcal{B}}\mathcal{B} + \sum_\lambda f_\lambda \hat{f}_\lambda + n_\lambda \hat{n_\lambda}\right)\right) \tag{S25}$$

$$\times \exp\left(-\frac{S\sigma^2}{2}\int \mathrm{d}t\mathrm{d}t' \mathcal{A}\mathcal{B} - S\sum_\lambda \int \mathrm{d}t f_\lambda n_\lambda\right) \tag{S26}$$

$$\times \exp\left(\sum_i \psi_i\right), \tag{S27}$$

where we have used the shorthand

$$\begin{aligned}\psi_i = \ln\Bigg[\int \mathcal{D}\hat{x}_i \exp\Bigg(i\int \mathrm{d}t\hat{x}_i\left(\frac{\dot{x}_i}{x_i} - 1 + x_i - \sum_\lambda \mathcal{S}_i^{(\lambda)}\hat{n}_\lambda\right) - i\int \mathrm{d}t x_i \sum_\lambda \mathcal{I}_i^{(\lambda)}\hat{f}_\lambda \\ - i\int \mathrm{d}t\mathrm{d}t'\left(x_i(t)\hat{\mathcal{A}}(t,t')x_i(t') + \hat{x}_i(t)\hat{\mathcal{B}}(t,t')\hat{x}_i(t')\right)\Bigg)\Bigg]\end{aligned} \tag{S28}$$

In particular, the probability of observing a given set of parameters $\mathcal{A}, \hat{\mathcal{A}}, \mathcal{B}, \hat{\mathcal{B}}, f_\lambda, \hat{f}_\lambda, n_\lambda, \hat{n}_\lambda$ can be obtained by integrating over all possible trajectories that are compatible with them. This gives

$$\begin{aligned}\mathbb{P}[\mathcal{A},\dots] \propto \exp\left(iS\int \mathrm{d}t\mathrm{d}t'\left(\hat{\mathcal{A}}\mathcal{A}+\hat{\mathcal{B}}\mathcal{B}+\sum_\lambda f_\lambda\hat{f}_\lambda+n_\lambda\hat{n}_\lambda\right)\right)\\ \times\exp\left(-\frac{S\sigma^2}{2}\int \mathrm{d}t\mathrm{d}t'\mathcal{A}\mathcal{B}-S\sum_\lambda\int \mathrm{d}t f_\lambda n_\lambda\right)\\ \prod_i\left(\int \mathcal{D}x_i\exp(\psi_i)\right),\end{aligned} \tag{S29}$$

In the limit $S\to\infty$, all terms in the r.h.s. are exponential in $S$. This implies that the variables $\mathcal{A},\dots$ concentrate around the values that maximize $S^{-1}\ln\mathbb{P}[\mathcal{A},\dots]$. These can be obtained by differentiating w.r.t. hatted and un-hatted variables. For the latter we obtain the conditions

$$\frac{\partial}{\partial\mathcal{A}(t,t')}=i\hat{\mathcal{A}}(t,t')-\frac{\sigma^2}{2}\mathcal{B}(t,t')=0\Rightarrow\hat{\mathcal{A}}=-\frac{i\sigma^2}{2}\mathcal{B} \tag{S30}$$

$$\frac{\partial}{\partial\mathcal{B}(t,t')}=i\hat{\mathcal{B}}(t,t')-\frac{\sigma^2}{2}\mathcal{A}(t,t')=0\Rightarrow\hat{\mathcal{B}}=-\frac{i\sigma^2}{2}\mathcal{A} \tag{S31}$$

$$\frac{\partial}{\partial f_\lambda(t,t')}=i\hat{f}_\lambda(t,t')-n_\lambda(t,t')=0\Rightarrow\hat{f}_\lambda=n_\lambda \tag{S32}$$

$$\frac{\partial}{\partial n_\lambda(t,t')}=i\hat{n}_\lambda(t,t')-f_\lambda(t,t')=0\Rightarrow\hat{n}_\lambda=f_\lambda. \tag{S33}$$

For the hatted variables we get

$$\frac{\partial}{\partial\hat{\mathcal{A}}(t,t')}=i\mathcal{A}(t,t')-iS^{-1}\sum_j\langle x_j(t)x_j(t')\rangle_j=0\Rightarrow\mathcal{A}=S^{-1}\sum_j\langle x_j(t)x_j(t')\rangle_j \tag{S34}$$

$$\frac{\partial}{\partial\hat{\mathcal{B}}(t,t')}=i\mathcal{B}(t,t')-iS^{-1}\sum_j\langle \hat{x}_j(t)\hat{x}_j(t')\rangle_j=0\Rightarrow\mathcal{B}=S^{-1}\sum_j\langle \hat{x}_j(t)\hat{x}_j(t')\rangle_j \tag{S35}$$

$$\frac{\partial}{\partial\hat{f}_\lambda(t)}=if_\lambda(t)-iS^{-1}\sum_j\mathcal{I}_j^{(\lambda)}\langle x_j(t)\rangle_j=0\Rightarrow f_\lambda=S^{-1}\sum_j\mathcal{I}_j^{(\lambda)}\langle x_j\rangle_j \tag{S36}$$

$$\frac{\partial}{\partial\hat{n}_\lambda(t)}=in_\lambda(t)-iS^{-1}\sum_j\mathcal{S}_j^{(\lambda)}\langle \hat{x}_j(t)\rangle_j=0\Rightarrow n_\lambda=S^{-1}\sum_j\mathcal{S}_j^{(\lambda)}\langle \hat{x}_j\rangle_j, \tag{S37}$$

where for some functional $\mathcal{O}[x_j,\hat{x}_j]$ we have introduced the notation

$$\langle\mathcal{O}\rangle_j=\frac{\int\mathcal{D}\hat{x}_j\mathcal{D}x_j\mathcal{O}\exp\left(i\int\mathrm{d}t\left(\hat{x}_j\left(\frac{\dot{x}_j}{x_j}-1+x_j-\sum_\lambda\mathcal{S}_j^{(\lambda)}\hat{n}_\lambda\right)-x_j\sum_\lambda\mathcal{I}_j^{(\lambda)}\hat{f}_\lambda-i\int\mathrm{d}t'\left(x_j(t)\hat{\mathcal{A}}(t,t')x_j(t')+\hat{x}_j(t)\hat{\mathcal{B}}(t,t')\hat{x}_j(t')\right)\right)\right)}{\int\mathcal{D}\hat{x}_j\mathcal{D}x_j\exp\left(i\int\mathrm{d}t\left(\hat{x}_j\left(\frac{\dot{x}_j}{x_j}-1+x_j-\sum_\lambda\mathcal{S}_j^{(\lambda)}\hat{n}_\lambda\right)-x_j\sum_\lambda\mathcal{I}_j^{(\lambda)}\hat{f}_\lambda-i\int\mathrm{d}t'\left(x_j(t)\hat{\mathcal{A}}(t,t')x_j(t')+\hat{x}_j(t)\hat{\mathcal{B}}(t,t')\hat{x}_j(t')\right)\right)\right)} \tag{S38}$$

In particular one has

$$\left\langle \hat{x}_j(t)\hat{x}_j(t')\right\rangle_j = \left\langle \hat{x}_j(t)\right\rangle_j = 0 \Rightarrow \mathcal{B} = n_\lambda = 0, \tag{S39}$$

and therefore, plugging in the values obtained from the saddle-point conditions

$$\langle \mathcal{O}\rangle_j = \frac{\int \mathcal{D}\hat{x}_j \mathcal{O}[x_j,\hat{x}_j] \exp\left(i\int \mathrm{d}t\hat{x}_j\left(\frac{\dot{x}_j}{x_j} - 1 + x_j - \sum_\lambda \mathcal{S}_j^{(\lambda)} f_\lambda\right) - \frac{\sigma^2}{2}\int \mathrm{d}t\mathrm{d}t'\hat{x}_j(t)\mathcal{A}(t,t')\hat{x}_j(t')\right)}{\int \mathcal{D}\hat{x}_j \exp\left(i\int \mathrm{d}t\hat{x}_j\left(\frac{\dot{x}_j}{x_j} - 1 + x_j - \sum_\lambda \mathcal{S}_j^{(\lambda)} f_\lambda\right) - \frac{\sigma^2}{2}\int \mathrm{d}t\mathrm{d}t'\hat{x}_j(t)\mathcal{A}(t,t')\hat{x}_j(t')\right)}. \tag{S40}$$

Plugging in these values in eq. (S27) we obtain the following simplified expression for the probability of a trajectory:

$$\mathbb{P}\left[\vec{x}(t)\right] \propto \prod_j \int \mathcal{D}\hat{x}_i \exp\left(i\int \mathrm{d}t\hat{x}_j\left(\frac{\dot{x}_j}{x_j} - 1 + x_j - \sum_\lambda \mathcal{S}_j^{(\lambda)} f_\lambda\right) - \frac{\sigma^2}{2}\int \mathrm{d}t\mathrm{d}t'\hat{x}_j(t)\mathcal{A}(t,t')\hat{x}_j(t')\right). \tag{S41}$$

In particular the averages defined in eq. (S40) are in fact averages with respect to the probability measure induced by the trajectories themselves, i.e. averages like those defining $f_\lambda$ and $\mathcal{A}$ are to be self-consistently obtained from the effective dynamics that we are deriving with DMFT.

Because eq. (S41) appears as a product over all species, it follows that each species undergoes stochastic dynamics independent from the rest. Moreover, focusing on species $j$, the probability corresponds to the following stochastic process

$$\dot{x}_j = x_j\left(1 - x_j + \sum_\lambda \mathcal{S}_j^{(\lambda)} f_\lambda + \sigma\zeta_j(t)\right), \tag{S42}$$

where $\zeta_j$ is a colored Gaussian process with correlation $\langle \zeta_j(t)\zeta_j(t')\rangle = \mathcal{A}(t,t')$. The dynamics are now simplified to a set of independent one-dimensional stochastic processes. Yet, they are not fully solved since the variables $\mathcal{A}(t,t')$ and $f_\lambda(t)$ are self-consistently defined from the stochastic processes themselves. Solving this loop analytically is not possible.

### E Relationship between different types of averages

Throughout our calculation we will encounter three kinds of averages:

1. with respect to the randomness in the interaction coefficients.
2. with respect to the effective stochastic process, denoted by $\langle\cdot\rangle$.
3. with respect to species, denoted by $\overline{\cdot}$.

Because the stochastic process is an effective representation of the randomness in the interaction, averages w.r.t. it imply averages over the randomness in the interaction coefficients. If the initial abundances of the system are chosen randomly, then the average over the stochastic process also includes that kind of randomness. Otherwise, averages w.r.t. the stochastic process are equivalent to averages w.r.t. the interactions. This is a crucial ingredient of DMFT. Moreover, in situations where memory of the initial conditions is lost (such as in the Unique Fixed Point phase), averages this equivalence is true regardless of how initial conditions have been chose (randomly or not).

On the other hand, averages w.r.t. species are done for a given realization of the system, i.e. for a given realization of the randomness in the coefficients, and involve expressions of the form

$$\overline{f} = \frac{1}{S}\sum_i f_i, \tag{S43}$$

where $f_i$ is any function of species $i$. These are in principle different from averages w.r.t. interactions/stochastic processes. However, because the stochastic processes of different species are independent, the central limit theorem ensures that

$$\overline{f} = \frac{1}{S}\sum_i f_i = \frac{1}{S}\sum_i \langle f_i \rangle = \overline{\langle f \rangle}. \tag{S44}$$

In other words, averages over species imply averages over the stochastic process. In unstructured ecosystems, the statistical equivalence of species implies that the converse is also true. In our case, structure makes species different, so that the equivalence no longer holds.

#### F Obtaining the equilibrium equations

We now show how to go from the effective process eq. (S42) to the equilibrium equations. Under the assumption that the system indeed reaches a fixed point, the aforementioned loop can be solved mostly analytically.

Suppose that the system admits a fixed point with abundances $\{x_j^\star\}$. From the definition of $\mathcal{A}$,

$$\mathcal{A}(t,t') \xrightarrow[t,t'\to\infty]{} C^\star \equiv S^{-1}\sum_j \left\langle \left(x_j^\star\right)^2 \right\rangle_j = \overline{(x^\star)^2}, \tag{S45}$$

where the overline now indicates an average over the species. Note that because the processes are independent across species, averaging over species implies averaging over the dynamics of each species. Similarly,

$$f_\lambda(t) \xrightarrow[t\to\infty]{} f^\star_\lambda \equiv S^{-1}\sum_j \mathcal{I}^{(\lambda)}_j \left\langle x^\star_j \right\rangle_j = \overline{\mathcal{I}^{(\lambda)} x^\star}. \tag{S46}$$

Since $\mathcal{A}(t,t')$ is the correlator of the processes $\zeta_j$, its convergence implies that all the processes $\zeta_j$ converge at infinity to some (random) value $\zeta^\star_j$. Let $\xi^\star_j = \zeta^\infty_j/\sqrt{C^\star}$ . Then equating to zero the derivative in eq. (S42) gives

$$x^\star_j = 1 + \sum_\lambda \mathcal{S}^{(\lambda)}_j f^\star_\lambda + \gamma\xi^\star_j \text{ or } x^\star_j = 0. \tag{S47}$$

We recall that $\gamma = \sigma\sqrt{C^\star}$. The first solution is only feasible if it is positive. On the other hand, it can be easily checked that the second solution is unstable as soon as the first one is positive. Hence the general solution is

$$x^\star_j = \max\left(0, 1 + \sum_\lambda \mathcal{S}^{(\lambda)}_j f^\star_\lambda + \gamma\xi^\star_j\right), \tag{S48}$$

In particular, $\xi^\star_j$ is a Gaussian random variable with zero mean and unit variance.

Now that the law of each species's equilibrium value is obtained as a function of $f^\star_\lambda$ and $\gamma$, we obtain the self-consistent equations for these two variables. By definition

$$f^\star_\lambda = \overline{\mathcal{I}^{(\lambda)} x^\star} = \overline{\int \mathcal{D}z \max\left(0, 1 + \sum_\lambda \mathcal{S}^{(\lambda)} f^\star_\lambda + \gamma z\right)} = \gamma\overline{\mathcal{I}^{(\lambda)} \int \mathcal{D}z \max\left(0, x_+/\gamma + z\right)} \tag{S49}$$

$$C^\star = \overline{(x^\star)^2} = \overline{\int \mathcal{D}z \max\left(0, 1 + \sum_\lambda \mathcal{S}^{(\lambda)} f^\star_\lambda + \gamma z\right)^2} = \gamma^2 \overline{\int \mathcal{D}z \max\left(0, x_+/\gamma + z\right)^2}, \tag{S50}$$

where $x^+, \gamma$ are defined in the main text and $\mathcal{D}z$ now represents the probability measure of a standard Gaussian variable. These are the equations given in the main text up to the definition of $\omega_1(\cdot)$ and $\omega_2(\cdot)$.

#### G DMFT with non-uniform variances in the random part

We now briefly describe how to extend the previous calculation to the case where the random part of the interaction coefficients has a non-uniform variance. In other words, whenever $\langle z^2_{ij}\rangle = \sigma^2_{ij}$. Our calculation can be straightforwardly extended to accomodate virtually any $\sigma_{ij}$ pattern, but for the reasons stated in the main text this generality is not our goal here. We limit ourselves to the case of a rank-one standard deviation matrix that can be written $\sigma_{ij} = \sigma\kappa_i\nu_j$.

The relevance of this case is explained in the main text. In short, it arises whenever carrying capacities are not uniform and a rescaling by the carrying capacities is performed. In that case, the interaction coefficients are rescaled from $A_{ij}$ to $A_{ij}K_j/K_i$, leading to a rescaling of the strength of hidden interactions from $\sigma_{ij} = \sigma$ to $\sigma_{ij} = \sigma K_j/K_i$. Hence the effective equations for this case can be obtained by taking $\kappa_i = 1/K_i$ and $\nu_j = K_j$ in what follows.

The main difference in the DMFT calculation comes from the integral w.r.t. interaction coefficients eq. (S14), which now reads

$$\int \prod_{ij} \mathrm{d}z_{ij} \mathbb{P}\left[z_{ij}\right] \exp\left(i \sum_{ij} \int \frac{\sigma z_{ij}}{\sqrt{S}} \mathrm{d}t \hat{x}_i(t) x_j(t)\right) = \exp\left[-\frac{S\sigma^2}{2} \int \mathrm{d}t\mathrm{d}t' \left(\frac{1}{S}\sum_i \kappa_i^2 \hat{x}_i(t)\hat{x}_i(t')\right)\left(\frac{1}{S}\sum_i \nu_i^2 x_i(t) x_i(t')\right)\right]. \tag{S51}$$

This implies a straightforward modification of the parameters $\mathcal{A}$ and $\mathcal{B}$, and the rest of the calculation is essentially unchanged. In the end, the effective stochastic process is

$$\dot{x}_j = x_j \left(1 - x_j + \sum_\lambda \mathcal{S}_j^{(\lambda)} f_\lambda + \sigma \kappa_j \zeta_j(t)\right), \tag{S52}$$

and the correlator of $\zeta_i$ is now $C_\nu(t,t') \equiv \langle \zeta_i(t)\zeta_i(t')\rangle = \overline{\nu^2 x(t) x(t')}$. In particular the equilibrium distribution is changed to

$$x_j^\star = \max\left(0, 1 + \sum_\lambda \mathcal{S}_j^{(\lambda)} f_\lambda^\star + \gamma_\nu \kappa_j \xi_j^\star\right), \tag{S53}$$

where we have defined $\gamma_\nu = \sigma\sqrt{C_\nu^\star}$. The fixed point equations read

$$f_\lambda^\star = \overline{\mathcal{I}^{(\lambda)} x^\star} = \overline{\mathcal{I}^{(\lambda)} \int \mathcal{D}z \max\left(0, 1 + \sum_\lambda \mathcal{S}^{(\lambda)} f_\lambda^\star + \kappa\gamma_\nu z\right)} = \gamma_\nu\, \overline{\kappa \mathcal{I}^{(\lambda)} \omega_1(x^+/\kappa\gamma_\nu)} \tag{S54}$$

$$C_\nu^\star = \overline{\nu^2 \left(x^\star\right)^2} = \overline{\nu^2 \int \mathcal{D}z \max\left(0, 1 + \sum_\lambda \mathcal{S}^{(\lambda)} f_\lambda^\star + \kappa\gamma_\nu z\right)^2} = \gamma_\nu^2\, \overline{\nu^2 \kappa^2 \omega_2(x^+/\kappa\gamma_\nu)}. \tag{S55}$$

#### H Obtaining the stability conditions

The stability can be analyzed by adding an initial perturbation to the abundances of the fixed point and checking whether the system returns to the fixed point or not. For small perturbations, we do so by linearizing the effective dynamics around the fixed point. This is therefore equivalent to checking the stability of the

Jacobian. The linearized stochastic differential equation reads

$$\frac{\mathrm{d}}{\mathrm{d}t}\delta x_j(t) = \delta x_j(t)\left(1 - x_j^\star + \sum_\lambda \mathcal{S}_j^{(\lambda)} f_\lambda^\star + \gamma\xi_j^\star\right) + x_j^\star\left(-\delta x_j(t) + \sum_\lambda \mathcal{S}_j^{(\lambda)}\delta f_\lambda(t) + \sigma\delta\zeta_j(t)\right), \tag{S56}$$

where we use the notation $\delta_j x(t) = x_j(t) - x_j^\star$. The fluctuations $\delta f_\lambda$ and $\delta\xi_j$ are self-consistently induced by the fluctuations in the $\delta x_j(t)$'s, just like $f_\lambda$ and $\xi_j$ are induced by the $x_j$'s. As such they have to obey the relations

$$\delta f_\lambda(t) = \overline{\mathcal{I}^{(\lambda)}\delta x(t)} \tag{S57}$$

$$\langle\xi_j(t)\xi_j(t')\rangle = \overline{\delta x(t)\delta x(t')}. \tag{S58}$$

The first term in the r.h.s. of eq. (S56) is zero for extant species, due to eq. (S48). It is negative for extinct species for the same reason. As such, it can never cause an instability. The second term is zero for extinct species and non-zero for extant species. As such, only the second term can cause the instability. We are therefore left to study the following stochastic process *over the set of extant species*

$$\frac{\mathrm{d}}{\mathrm{d}t}\delta x_j(t) = x_j^\star\left(-\delta x_j(t) + \sum_\lambda \mathcal{S}_j^{(\lambda)}\delta f_\lambda(t) + \sigma\delta\xi_j(t)\right). \tag{S59}$$

Given that the $\xi_j$'s are Gaussian processes, the $\delta\xi_j$'s are also Gaussian processes. We can therefore fully characterize themselves through their mean and two-point correlations. In particular, taking the mean and using eq. (S58) gives

$$\frac{\mathrm{d}}{\mathrm{d}t}\langle\delta x_j(t)\rangle = x_j^\star\left(-\langle\delta x_j(t)\rangle + \sum_{\lambda,i}\mathcal{S}_j^{(\lambda)}\mathcal{I}_i^{(\lambda)}\langle\delta x_i(t)\rangle\right) \Rightarrow \frac{\mathrm{d}}{\mathrm{d}t}\left\langle\vec{\delta x}\right\rangle = \mathcal{J}\left\langle\vec{\delta x}\right\rangle, \tag{S60}$$

where the matrix $\mathcal{J}$ is defined as

$$\mathcal{J}_{ij} = -x_i^\star\left(\delta_{ij} - \frac{\mu_{ij}}{S}\right). \tag{S61}$$

Positive eigenvalues of $\mathcal{J}$ indicate an instability of the system. Note that it makes no difference to eigenvalues whether we include or exclude the extinct species in the definition of this matrix. As such, we include them to be able to work with our knowledge of the full structure matrix. This equation can be used to predict the stability of the fixed points of the system presented in Sec. 3.3. In particular, Fig. S1 shows the spectrum of $\mathcal{J}$ for the two central rows of Fig. 5, that is right before and right after the stabilization of the fixed point.

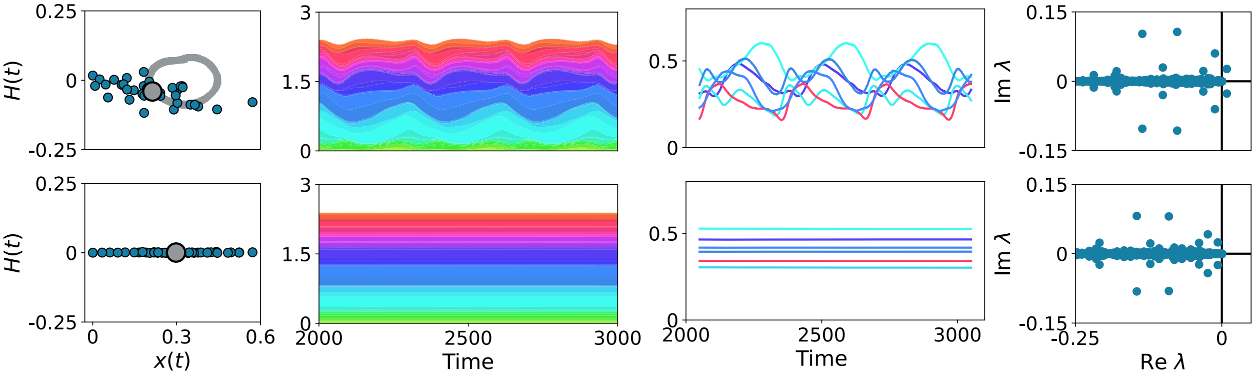


FIG. S1: SPECTRUM OF THE PSEUDO-JACOBIAN BEFORE AND AFTER STABILIZATION. We consider here the same system as in Fig. 5 for $\sigma = 1$ and $\sigma = 1.75$ (panels B and C), that is just before and just after the equilibrium is stabilized. A pair of complex outliers is seen to signal the transition from macroscopic cycles to stability.

Before stabilization, a pair of complex outliers has positive real part: thus the system is unstable and, in fact, displays cyclic oscillations. As the complex outliers get closer to the real axis, a Hopf bifurcation takes place: the cycles disappear and the fixed point becomes stable. We now turn our attention to the instability condition associated to the correlations. For this one, it is convenient to take the Laplace transform of the linearized effective process. This yields

$$s\hat{\delta x}_j(s) = x_j^\star \left(-\hat{\delta x}_j(s) + \sum_\lambda \mathcal{S}_j^{(\lambda)} \hat{\delta f}_\lambda(s) + \sigma \hat{\delta\xi}_j(s)\right) + \delta x_j(0), \tag{S62}$$

where $\delta x_j(0)$ is the initial perturbation to the abundance of species $j$. Given the Final Value Theorem for Laplace transforms, stability requires that

1. $\hat{\delta x}_j(s)$ has no poles on the right half of the complex plane.

2. $s\hat{\delta x}_j(s) \to 0$ when $s \to 0$.

Without loss of generality, we can assume that we deal with a set of parameters for which the mean is stable in the sense of eq. (S60). In that case $\left\langle s\hat{\delta x}_j(s)\right\rangle \to 0$ as $s \to 0$ and therefore $s\hat{\delta f}_\lambda(s) \to 0$ as well. Now multiply by $s$, take squared moduli and average in eq. (S62),

$$\left|\frac{s}{x_j^\star}+1\right|^2 \left\langle \left|s\hat{\delta x}_j(s)\right|^2\right\rangle = \left|\sum_\lambda \mathcal{S}_j^{(\lambda)} s\hat{\delta f}_\lambda(s)\right|^2 + \sigma^2\left\langle\left|s\hat{\delta\xi}_j(s)\right|^2\right\rangle + |s|^2\delta x_j(0)^2. \tag{S63}$$

The loop is closed using the following self-consistency condition between the $\hat{\delta x}_j(s)$'s and the $\hat{\delta \xi}_j(s)$'s that stems from eq. (S58)

$$\left\langle \left| \hat{\delta \xi}_j(s) \right|^2 \right\rangle = S^{-1} \sum_j \left\langle \left| \hat{\delta x}_j(s) \right|^2 \right\rangle = \overline{\left| \hat{\delta x}(s) \right|^2}. \tag{S64}$$

As usual, the overline denotes an average over species, which in turn implies also the average $\langle\rangle$ over the effective process. Averaging over species in eq. (S63) gives

$$\overline{\left| s\hat{\delta x}(s) \right|^2} = \frac{\left| \sum_\lambda \mathcal{S}_j^{(\lambda)} s \hat{\delta f}_\lambda(s) \right|^2 + |s|^2 \overline{\delta x(0)^2}}{\left| 1 + \frac{s}{x^\star} \right|^2 - \sigma^2 \phi}. \tag{S65}$$

The fraction of extant species $\phi$ appears because the average over species includes both extant and extinct species, but extinct species have a null contribution. Now take the limit $s \to 0$. By assumption, the first term in the r.h.s. of eq. (S63) vanishes and we therefore have

$$\overline{\left| s\hat{\delta x}(s) \right|^2} \sim \frac{|s|^2 \overline{\delta x(0)^2}}{1 - \sigma^2 \phi} \to 0, \tag{S66}$$

which is vanishing, unless the condition $1 = \sigma^2 \phi$ is met. This signals the instability. For $\sigma^2 \phi > 1$ the condition is not met, but the function has a pole on the right half of the complex plane, so there is a divergence nonetheless. For $\sigma^2 \phi < 1$ the expression goes to zero and has no poles on the right half of the complex plane, so there is no divergence.

Note that eq. (S66) was obtained by taking averages over species, so one could wonder whether instabilities could happen for individual species that are not captured by this analysis. This is not the case given eq. (S63). Indeed,

$$\left\langle \left| s\hat{\delta x}_j(s) \right|^2 \right\rangle \sim_{s \to 0} \frac{\sigma^2 |s|^2 \overline{\delta x(0)^2}}{1 - \sigma^2 \phi} + |s|^2 \delta x_j(0)^2, \tag{S67}$$

and therefore the same analysis applies to each individual species. In other words, this divergence occurs for all species at the same point.

#### I Illustrations for particular structures

As explained in the main text, different hypothesis on the functions, in small number $n_F$, that underpin species interaction at the community level produce different structural matrices. These provide the low-dimensional 'skeleton' that gets blurred – and never cancels out – when randomness is added.

Figure S2 provides a schematic representation of the functional traits that define the structure in two cases of

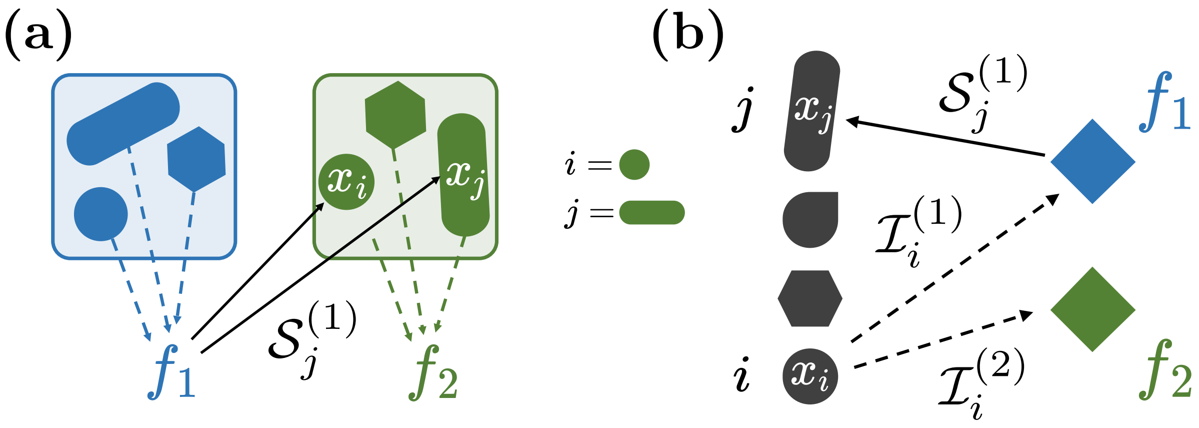


Fig. S2: Examples of ecosystem structure and their formalization. **A** The community is divided in two functional groups, whose total abundances $f_1, f_2$ are the relevant collective functions. Species in group $\lambda \in \{1,2\}$ contribute to the abundance of their group with equal impact traits, $\mathcal{I}_i^{(\lambda)} = 1$. Conversely, a group acts upon the rest of the community through its total biomass $f_\lambda$, whose effect on any species is weighted by the sensitivity traits $\mathcal{S}_i^{(\lambda)}$ (shown only for the effect of the blue group on two species of the green group). Here, only interactions between functional groups are measurable, and community dynamics is hence described in terms of the total abundance of species belonging to each group - while within-group variability is ignored. In this formalization, disordered gLVEs [3] – where the total biomass is the only collective function – corresponds to the existence of a single group. **B** Interactions between species are mediated by two resources of concentrations $f_1, f_2$, which provide the collective functions. Each species affects the function by either consuming the resource or producing a resource, as encoded by the impact traits $\mathcal{I}^{(\lambda)}$ (dotted lines, shown only for species $i$). Conversely, it is affected by these resources through the sensitivity traits $\mathcal{S}^{(\lambda)}$ (shown only for species $j$). This scenario reflects the classic assumptions of consumer-resource models, where impact traits are the consumption rates, and sensitivity traits are the rate of transformation into units of species abundance [4, 5].

ecological relevance: species belong to (homogeneous) functional groups (panel a) and a consumer-resource model (panel b).

Below, we detail three examples of structure that are mentioned in the main text.

##### I.1 Example of a structure that produces nontrivial species abundance distributions.

Figure S3 illustrates the fact that when species traits are drawn from a broader distribution than the typical scale $\gamma$ arising from disorder, arbitrary Species Abundance Distributions can be observed, such as power-law distributions (as well as multimodal ones). This reflect the fact that the structured component of the interactions differentiates species more than the disordered one, even though it is not necessarily larger in total magnitude.

##### I.2 Example of a structure that has no effect on statistical patterns

We consider now a toy model where interactions are underpinned by neutrally evolving traits. The distance $d_{ij}$ of two species along a phylogenetic tree along which traits diverge then determines the intensity of

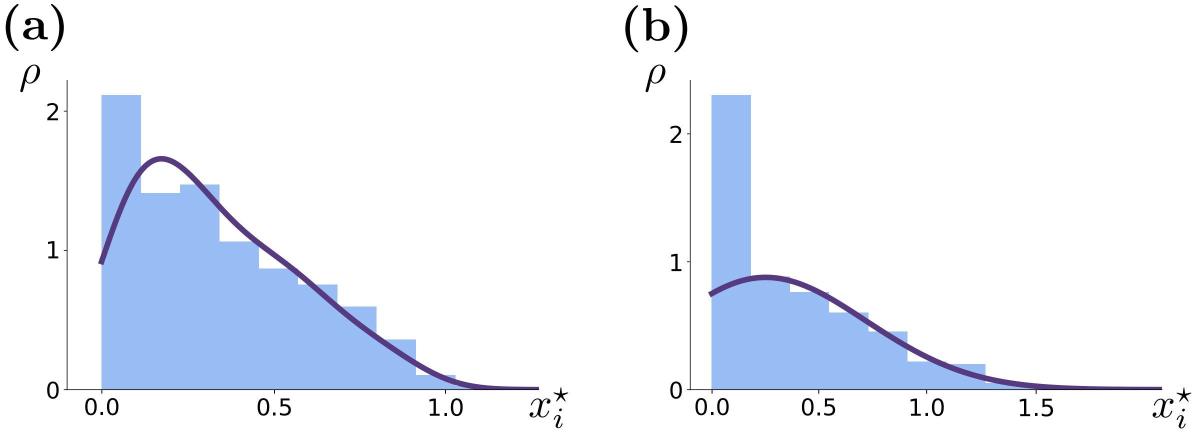


FIG. S3: Comparison of the SAD of a simulated community with $S = 1000$ (histograms) and the theoretical predictions of eq. (13) (lines) for two different levels of random variation in the interactions (**A**, $\sigma = 0.25$ and **B** , $\sigma = 0.75$). Species abundances are affected by a single function: the total biomass (meaning that they all have the same impact trait $\mathcal{I}_j^{(\lambda)} = 1$). However, the sensitivity trait $\mathcal{S}_i^{(\lambda)}$ of a species $i$ is drawn from a power-law distribution with exponent $\alpha = 2$ on the interval $[0, 3]$. The SAD that reflects the power-law distribution of the traits, discernible for small $\sigma$, turns into a truncated Gaussian as the magnitude of disorder increases, progressively blurring the structural features. The shape of the SAD is well predicted from the sole values of $\alpha$ and $\sigma$, even if the exact functional traits of each species in the community are unknown.

interactions $\mu_{ij} = f(d_{ij})$. The simplest form of such phylogeny is a balanced binary tree, which produces following structure for the interaction matrix

$$\begin{pmatrix} 0 & f(1) & f(2) & f(2) & f(3) & f(3) & f(3) & f(3) \\ f(1) & 0 & f(2) & f(2) & f(3) & f(3) & f(3) & f(3) \\ f(2) & f(2) & 0 & f(1) & f(3) & f(3) & f(3) & f(3) \\ f(2) & f(2) & f(1) & 0 & f(3) & f(3) & f(3) & f(3) \\ f(3) & f(3) & f(3) & f(3) & 0 & f(1) & f(2) & f(2) \\ f(3) & f(3) & f(3) & f(3) & f(1) & 0 & f(2) & f(2) \\ f(3) & f(3) & f(3) & f(3) & f(2) & f(2) & 0 & f(1) \\ f(3) & f(3) & f(3) & f(3) & f(2) & f(2) & f(1) & 0 \end{pmatrix} \tag{S68}$$

corresponding to the nested structural matrix represented in Fig. S4 a. The addition of randomness partially blurs the structure (Fig. S4 b), so that one may expect that it affects the community statistics.

The structural matrix has the interesting symmetry that $(1, \ldots, 1)$ is an eigenvector. This implies the existence of a fixed point that is symmetric under permutation of species, i.e. such that the distribution of abundances is the same for all species. In particular, the equilibrium equations at this fixed point are the same as those of the purely disordered gLV with an effective mean interaction

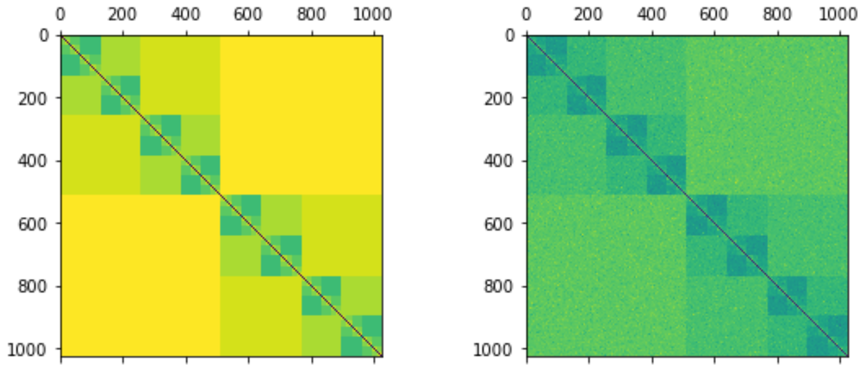


FIG. S4: Interaction matrix structured by binary phylogenetic tree with $S = 1024$ species. **Left:** Deterministic structure. **Right:** Superposition of structure and disorder.

$$\mu^{\text{eff}} = \sum_d 2^{d-1} f(d).$$

Hence, given that the macroscopic equations only depend on the abundances at equilibrium, the phase diagram of such system coincides with that of the purely disordered gLV. Therefore, randomness in this case does not change the qualitative properties of the community dynamics.

##### I.3 Model with a high number of groups

We consider a case when the number of groups, or functions, is large (but small relative to the total number of species). The gLV equations can in this case exhibit out-of equilibrium dynamics. Following [6], the group-level interaction matrix $M$ of size $n_F \times n_F$ is built by sampling all its coefficients randomly and independently from a normal distribution of mean $\mu_g$ and variance $\Sigma^2$. In particular, the mean and variance are of order one and do not scale with $n_F$. For $n_F$ large enough (we take $n_F = 150$ as our reference case), the $n_F$-dimensional gLV with interaction matrix $M$ (and a small migration) exhibits turnover dynamics, whereby at any point in time the number of dominant species in the community is small but the identity of such species changes over time due to invasions and extinctions[6].

We add randomness by dividing each of the $n_F$ groups into a large number $s$ species for a total of $S = n_F \times s$ species (in the main text we use $s = 40$ and $S = 6000$) and letting the structural matrix be dictated by group affiliation: if species $i, j$ belong respectively to groups $a, b$ then $\mu_{ij} = M_{ab}/s$. Note that in going from the macroscopic to the microscopic model we divide by the number of species per group, so that the interactions at the species level are weak, but the aggregated interactions at the group level are strong as in the case where groups are represented by only one species.

For $\sigma = 0$, species within groups are identical and synchronize, so that the long-term trajectory of each group is the same as that of the species that belong to it. In particular, both species and groups follow chaotic trajectories dictated by turnover at the group level. Upon increasing $\sigma$, species within groups cease to be identical and a variety of dynamical regimes is observed, as depicted in Fig. 5.
